## Supplemental figures and tables for "*PTC2* region genotypes counteract *Biomphalaria glabrata* population differences between M-line and BS90 in resistance to infection by *Schistosoma mansoni*"

Supplemental Figure S1. Summary of the populations created and samples taken for analysis in this study.

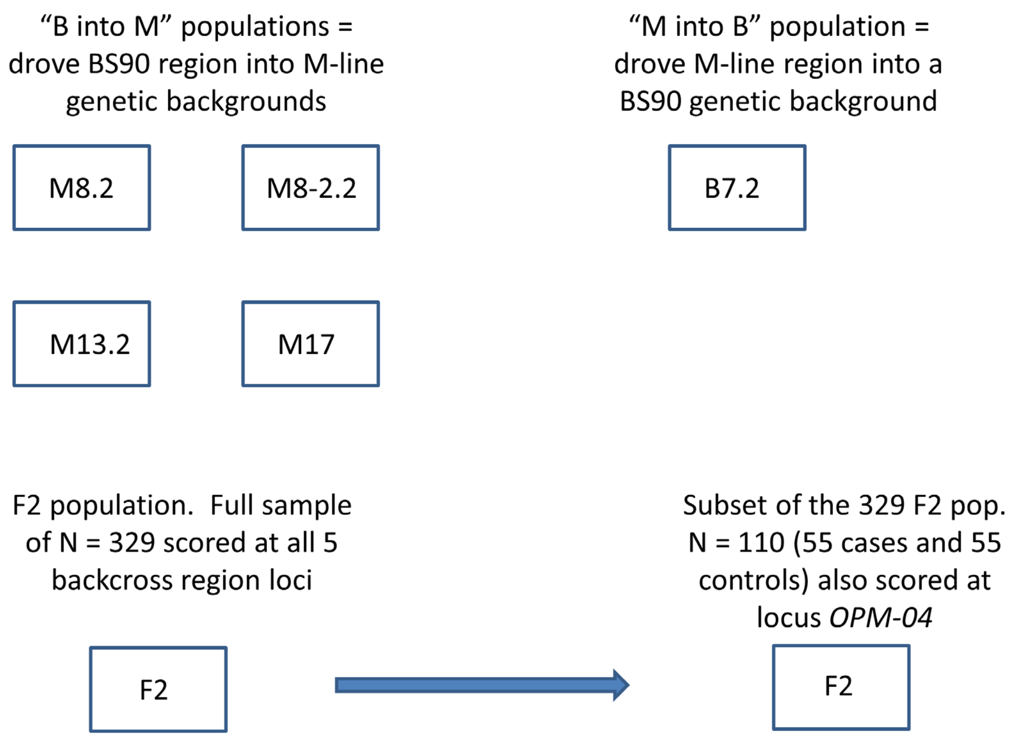

Supplemental Figure S2. *OPM-04* locus structure and sequence

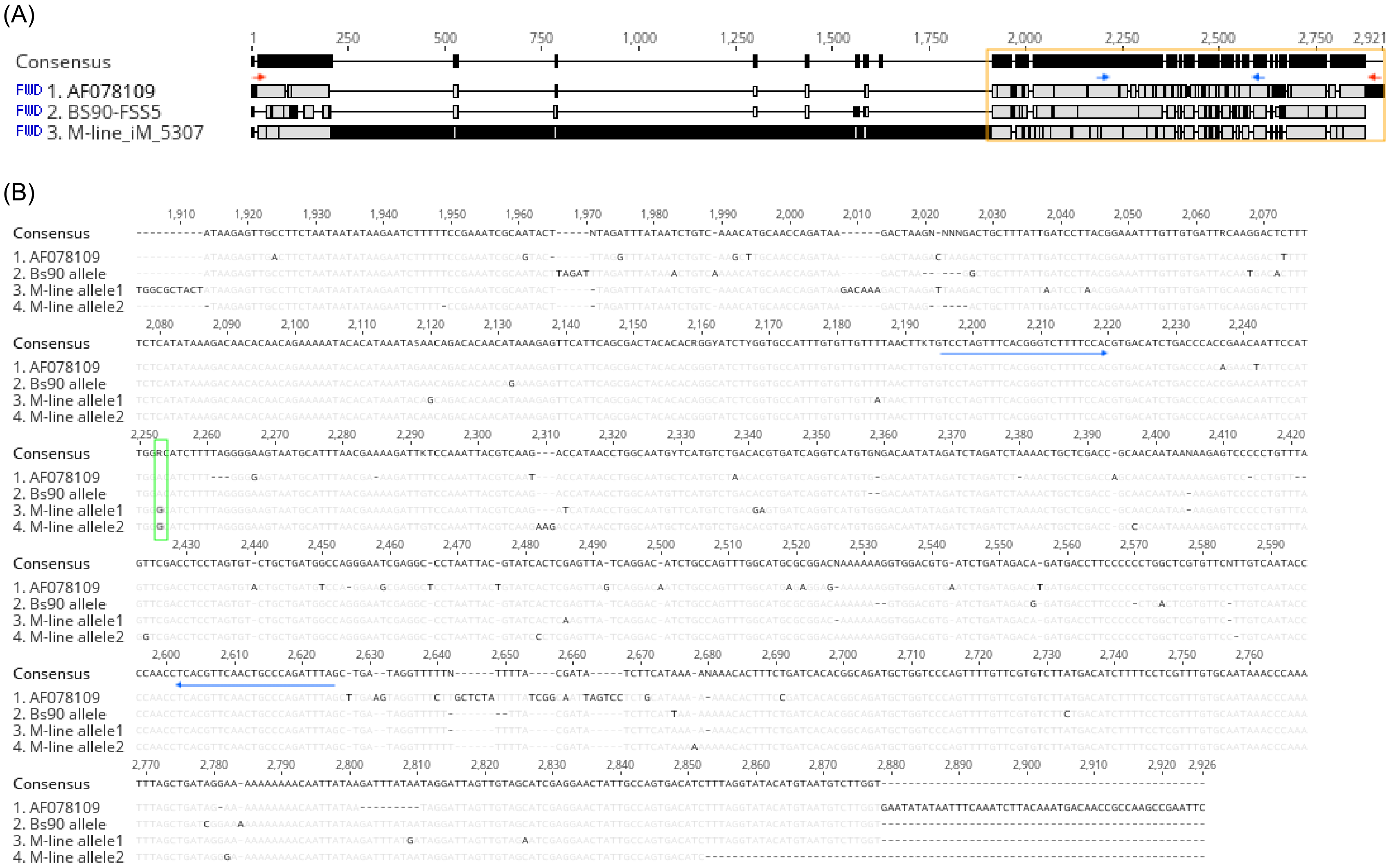

Table S1. Primer sequences (5' to 3') and amplicon sizes.

| Locus | Primer Forward | Primer Reverse | amplicon size BS90 | amplicon size M-line |
|---|---|---|---|---|
| *up2* | AGTAACAACAGTAAACGTAACTTTCT | CCAGTACTGCGATTGGTTAGGGG | 928 | 503 |
| *up1* | TCATTTATCCCTTAGTCTGGTGG | CAAGACCCATGCGATCACAG | 880 | 485 |
| *0* | ATAGAATGATATAGTGGAGCTTGCA | GATCCAGGAATTTGTTTCGGAAGTA | 751 | 979 |
| *dn1* | GGCAGACAATCTTGAGACAGTAG | CTAGTCAAGAACAAGAGCGAAAG | 596 | 213 |
| *dn2* | CATGCAATAACTAACCTGATGCCAA | GTAACTGATCACGAGAGGGAAAACC | 1284 | 1086 |
| *OPM-04* | GTCCTAGTTTCACGGGTCTTTTCCAC | CTAAATCTGGGCAGTTGAACGTGAG | 413 | 417/422 |

Table S2. Raw genotype and infection data used for logistic regressions and figures.

"B into M" populations, locus *PTC2*

| population | *PTC2* genotype | infected?(1=yes) | number observed |
|---|---|---|---|
| M8.2 | BB | 1 | 8 |
| | BB | 0 | 14 |
| | BM | 1 | 13 |
| | BM | 0 | 79 |
| | MM | 1 | 5 |
| | MM | 0 | 40 |
| M8-2.2 | BB | 1 | 15 |
| | BB | 0 | 4 |
| | BM | 1 | 31 |
| | BM | 0 | 19 |
| | MM | 1 | 9 |
| | MM | 0 | 11 |
| M13.2 | BB | 1 | 10 |
| | BB | 0 | 11 |
| | BM | 1 | 17 |
| | BM | 0 | 28 |
| | MM | 1 | 6 |
| | MM | 0 | 13 |
| M17 | BB | 1 | 8 |
| | BB | 0 | 4 |
| | BM | 1 | 30 |
| | BM | 0 | 11 |
| | MM | 1 | 11 |
| | MM | 0 | 4 |

"M into B" population, locus *PTC2*

| population | *PTC2* genotype | infected?(1=yes) | number observed |
|---|---|---|---|
| B7.2 | BB | 1 | 3 |
| B7.2 | BB | 0 | 30 |
| B7.2 | BM | 1 | 7 |
| B7.2 | BM | 0 | 57 |
| B7.2 | MM | 1 | 1 |
| B7.2 | MM | 0 | 18 |

F2 entire sample, N = 329, locus *PTC2*

| population | *PTC2* genotype | infected?(1=yes) | number observed |
|---|---|---|---|
| F2(N=329) | BB | 0 | 69 |
| | BB | 1 | 24 |
| | BM | 0 | 140 |
| | BM | 1 | 29 |
| | MM | 0 | 58 |
| | MM | 1 | 9 |

F2 cases & controls, locus *PTC2*

| population | *PTC2* genotype | infected?(1=yes) | number observed |
|---|---|---|---|
| F2(case/control) | BB | 0 | 10 |
| | BB | 1 | 23 |
| | BM | 0 | 36 |
| | BM | 1 | 24 |
| | MM | 0 | 9 |
| | MM | 1 | 7 |

F2 cases & controls, locus *OPM-04*

| population | *OPM-04* genotype | infected?(1=yes) | number observed |
|---|---|---|---|
| F2(case/control) | BB | 0 | 8 |
| | BB | 1 | 13 |
| | BM | 0 | 28 |
| | BM | 1 | 27 |
| | MM | 0 | 16 |
| | MM | 1 | 15 |

Table S3. Binary logistic regression output (obtained using SYSTAT 13.2).

**“B into M” populations, locus *0***

Additive model. AIC = 458.78

| Parameter Estimates | | | | | | |
|---|---|---|---|---|---|---|
| Parameter | Estimate | Standard Error | Z | p-Value | 95% Confidence Interval | |
| | | | | | Lower | Upper |
| CONSTANT | 0.810 | 0.397 | 2.041 | 0.041 | 0.032 | 1.587 |
| GENOTYPE | 0.790 | 0.352 | 2.246 | 0.025 | 0.101 | 1.480 |
| POP$_13.2 | -0.684 | 0.551 | -1.241 | 0.214 | -1.764 | 0.396 |
| POP$_17 | -1.597 | 0.650 | -2.457 | 0.014 | -2.871 | -0.323 |
| POP$_8-2.2 | -2.080 | 0.595 | -3.496 | 0.000 | -3.246 | -0.914 |
| POP$_13.2*GENOTYPE | -0.447 | 0.482 | -0.928 | 0.354 | -1.392 | 0.498 |
| POP$_17*GENOTYPE | -0.946 | 0.556 | -1.700 | 0.089 | -2.036 | 0.145 |
| POP$_8-2.2*GENOTYPE | -0.038 | 0.498 | -0.077 | 0.939 | -1.015 | 0.938 |

| Odds Ratio Estimates | | | | |
|---|---|---|---|---|
| Parameter | Odds Ratio | Standard Error | 95% Confidence Interval | |
| | | | Lower | Upper |
| GENOTYPE | 2.204 | 0.776 | 1.106 | 4.394 |
| POP$_13.2 | 0.505 | 0.278 | 0.171 | 1.486 |
| POP$_17 | 0.202 | 0.132 | 0.057 | 0.724 |
| POP$_8-2.2 | 0.125 | 0.074 | 0.039 | 0.401 |
| POP$_13.2*GENOTYPE | 0.639 | 0.308 | 0.249 | 1.645 |
| POP$_17*GENOTYPE | 0.388 | 0.216 | 0.131 | 1.156 |
| POP$_8-2.2*GENOTYPE | 0.962 | 0.479 | 0.362 | 2.555 |

M-allele dominant model. AIC = 459.61

| Parameter Estimates | | | | | | |
|---|---|---|---|---|---|---|
| Parameter | Estimate | Standard Error | Z | p-Value | 95% Confidence Interval | |
| | | | | | Lower | Upper |
| CONSTANT | 0.560 | 0.443 | 1.263 | 0.207 | -0.309 | 1.428 |
| POP$_13.2 | -0.464 | 0.622 | -0.746 | 0.456 | -1.684 | 0.756 |
| POP$_17 | -1.253 | 0.756 | -1.657 | 0.097 | -2.734 | 0.229 |
| POP$_8-2.2 | -1.881 | 0.716 | -2.626 | 0.009 | -3.285 | -0.477 |
| GENOTYPE | 1.329 | 0.510 | 2.605 | 0.009 | 0.329 | 2.329 |
| POP$_13.2*GENOTYPE | -0.846 | 0.721 | -1.175 | 0.240 | -2.259 | 0.566 |
| POP$_17*GENOTYPE | -1.642 | 0.852 | -1.926 | 0.054 | -3.312 | 0.029 |
| POP$_8-2.2*GENOTYPE | -0.295 | 0.797 | -0.370 | 0.711 | -1.857 | 1.267 |

| Odds Ratio Estimates | | | | |
|---|---|---|---|---|
| Parameter | Odds Ratio | Standard Error | 95% Confidence Interval | |
| | | | Lower | Upper |
| POP$_13.2 | 0.629 | 0.391 | 0.186 | 2.129 |
| POP$_17 | 0.286 | 0.216 | 0.065 | 1.257 |
| POP$_8-2.2 | 0.152 | 0.109 | 0.037 | 0.620 |
| GENOTYPE | 3.778 | 1.928 | 1.390 | 10.270 |
| POP$_13.2*GENOTYPE | 0.429 | 0.309 | 0.104 | 1.761 |
| POP$_17*GENOTYPE | 0.194 | 0.165 | 0.036 | 1.029 |

| Odds Ratio Estimates | | | | |
|---|---|---|---|---|
| Parameter | Odds Ratio | Standard Error | 95% Confidence Interval | |
| | | | Lower | Upper |
| POP$_8-2.2*GENOTYPE | 0.744 | 0.593 | 0.156 | 3.551 |

**F2 population, entire sample (N = 329), locus *up1***

Additive model. AIC = 318.17

| Parameter Estimates | | | | | | |
|---|---|---|---|---|---|---|
| Parameter | Estimate | Standard Error | Z | p-Value | 95% Confidence Interval | |
| | | | | | Lower | Upper |
| CONSTANT | 1.093 | 0.219 | 4.990 | 0.000 | 0.664 | 1.522 |
| GENOTYPE | 0.428 | 0.210 | 2.041 | 0.041 | 0.017 | 0.840 |

| Odds Ratio Estimates | | | | |
|---|---|---|---|---|
| Parameter | Odds Ratio | Standard Error | 95% Confidence Interval | |
| | | | Lower | Upper |
| GENOTYPE | 1.535 | 0.322 | 1.017 | 2.315 |

M-allele dominant model. AIC = 318.53

| Parameter Estimates | | | | | | |
|---|---|---|---|---|---|---|
| Parameter | Estimate | Standard Error | Z | p-Value | 95% Confidence Interval | |
| | | | | | Lower | Upper |
| CONSTANT | 1.056 | 0.237 | 4.456 | 0.000 | 0.592 | 1.521 |
| GENOTYPE | 0.595 | 0.296 | 2.010 | 0.044 | 0.015 | 1.174 |

| Odds Ratio Estimates | | | | |
|---|---|---|---|---|
| Parameter | Odds Ratio | Standard Error | 95% Confidence Interval | |
| | | | Lower | Upper |
| GENOTYPE | 1.812 | 0.536 | 1.015 | 3.236 |

**F2 population, 55 cases and 55 controls subset (N=110), locus *up1***

Additive model. AIC = 150.05

| Parameter Estimates | | | | | | |
|---|---|---|---|---|---|---|
| Parameter | Estimate | Standard Error | Z | p-Value | 95% Confidence Interval | |
| | | | | | Lower | Upper |
| CONSTANT | -0.552 | 0.326 | -1.697 | 0.090 | -1.190 | 0.086 |
| GENOTYPE | 0.678 | 0.310 | 2.186 | 0.029 | 0.070 | 1.287 |

| Odds Ratio Estimates | | | | |
|---|---|---|---|---|
| Parameter | Odds Ratio | Standard Error | 95% Confidence Interval | |
| | | | Lower | Upper |
| GENOTYPE | 1.971 | 0.612 | 1.073 | 3.621 |

M-allele dominant model. AIC = 147.25

| Parameter Estimates | | | | | |
|---|---|---|---|---|---|
| Parameter | Estimate | Standard Error | Z | p-Value | 95% Confidence Interval |

| | | | | | 95% CI Lower | Upper |
|---|---|---|---|---|---|---|
| | | | | | Lower | Upper |
| CONSTANT | -0.833 | 0.379 | -2.199 | 0.028 | -1.575 | -0.091 |
| MDOM | 1.206 | 0.445 | 2.710 | 0.007 | 0.334 | 2.078 |

**Odds Ratio Estimates**

| Parameter | Odds Ratio | Standard Error | 95% Confidence Interval | |
|---|---|---|---|---|
| | | | Lower | Upper |
| MDOM | 3.339 | 1.485 | 1.396 | 7.985 |

**F2 population, 55 cases and 55 controls subset (N=110), <u>locus *OPM-04*</u>**

Additive Model.  AIC = 151.09

**Parameter Estimates**

| Parameter | Estimate | Standard Error | Z | p-Value | 95% Confidence Interval | |
|---|---|---|---|---|---|---|
| | | | | | Lower | Upper |
| CONSTANT | -0.486 | 0.449 | -1.080 | 0.280 | -1.366 | 0.395 |
| GENOTYPE | 0.532 | 0.498 | 1.067 | 0.286 | -0.445 | 1.509 |

**Odds Ratio Estimates**

| Parameter | Odds Ratio | Standard Error | 95% Confidence Interval | |
|---|---|---|---|---|
| | | | Lower | Upper |
| GENOTYPE | 1.702 | 0.849 | 0.641 | 4.522 |

M-allele dominant model. AIC = 151.47

**Parameter Estimates**

| Parameter | Estimate | Standard Error | Z | p-Value | 95% Confidence Interval | |
|---|---|---|---|---|---|---|
| | | | | | Lower | Upper |
| CONSTANT | -0.327 | 0.366 | -0.895 | 0.371 | -1.044 | 0.389 |
| GENOTYPE | 0.247 | 0.282 | 0.877 | 0.380 | -0.306 | 0.801 |

**Odds Ratio Estimates**

| Parameter | Odds Ratio | Standard Error | 95% Confidence Interval | |
|---|---|---|---|---|
| | | | Lower | Upper |
| GENOTYPE | 1.281 | 0.361 | 0.737 | 2.227 |
